# Experimental population demography reveals sex specific density dependence as an outcome of sexual conflict

**DOI:** 10.64898/2026.02.09.704738

**Authors:** Anneli Brändén, Miguel A. Gomez-Llano, Stephen P. De Lisle

## Abstract

Many demographic models assume that only females matter for population dynamics. However, theory and evidence of sexual conflict suggest that males can affect female fitness through mating competition between and within the sexes, yet it is unclear how such effects may influence population dynamics. We used experimental population demography to understand how sexual conflict affects offspring recruitment in *Drosophila melanogaster*, a model species for studying the evolution of sexual conflict. By manipulating sex ratio and male/female density independently in a response surface design we found that increasing male density, and thereby the intensity of sexual conflict, led to fewer offspring per female, but that effect was nearly half the strength of female density dependence. Consistent with this, our best fitting birth function showed female dominance of births with sex-specific density dependence, indicating that males have a demographic effect even if females have demographic dominance. Our results confirm that females have a larger influence than males on offspring recruitment, however, more importantly our result increases our understanding about the demographic effects males have through sexual conflict.

## Introduction

Most demographic models are based on the assumption that only females matter for population growth (Caswell & Weeks, 1986). Female demographic dominance builds on the idea that only female characteristics limit fecundity and thus offspring production, while males just contribute sperm (Miller & Inouye, 2011). In addition to this, there is an assumption that only a few males per generation are required to provide sperm for all females in a population since sperm rarely is limiting (Eberhard, 1996; Parker, 1979). However, these assumptions are often violated in real populations as sexes often differ in life history (sexual maturity, fertility, life span and group dynamics) (Caswell & Weeks, 1986) and trait optima (food preferences, number of matings and habitats) (Chapman et al., 2003; Maklakov et al., 2008; Shine, 1989). Even though the primary sex ratio is often 1:1 in sexually reproducing animals (Miller, 1964; Papach et al., 2019; Rawls, 1913), the secondary sex ratio (operational sex ratio) can differ wildly between populations and species (Jenouvrier et al., 2010; Papach et al., 2019) due to higher mortality in one sex (Fisher, 1999) or differences in latency between mating events including females being unavailable to the mating pool when already occupied with some other reproductive stage (Arnqvist & Rowe, 2005; Darwin, 1872). Thus, a two-sex model may be far better to understand the population dynamics of natural populations of sexually reproducing organisms. A lingering empirical challenge is that we do not actually know what the best birth function (i.e., mathematical model describing birth rates based on female and male density in a two-sex population) is to describe the dynamics in real sexually reproducing populations (Miller & Inouye, 2011).

One reason for why the best birth function is unknown is the unclear role males play in population demography; do they matter or not? Males can influence demography in multiple ways but one common way is through sexual conflict, which can occur when males and females have different evolutionary interests (Arnqvist & Rowe, 2005; Parker, 1979). For example when the optimal number of matings is lower for females than for males (Burke & Bonduriansky, 2017; Eberhard, 1996), which can result in the evolution of male mating strategies that lower female fecundity. These costs of sexual conflict may not only affect individuals but also populations, as the costs of sexual conflict have been quantified and show to decrease female fecundity significantly (Gómez-Llano et al., 2024). This could mean that a higher proportion of males can increase the demographic costs of sexual conflict in a population (Krupa & Sih, 1993; Lauer et al., 1996), thus sex ratio is a vital component in describing population demography. As such, the connection between sexual conflict and population demography has been identified as a current unsolved challenge in understanding the role of sexual conflict in adaptation (Rowe & Rundle, 2021).

Another key element in population dynamics that can have important demographic effects is density (Kokko & Rankin, 2006). Studies on *Drosophila* using even sex ratios (e.g., Barker, 1973; Pearl, 1932; Wallace, 1974) have shown a clear connection between female density and offspring recruitment, but also a reduced per capita yield for females in larger populations. This means that population mean absolute fitness and an individual’s relative fitness can contradict each other in some cases, and points to the importance of competition and conflicts in the population (Darwin, 1872). Density also affects development, with high larval density resulting in a longer developmental time, smaller adults (Miller, 1964), greater variation in adult body size (Ashburner et al., 2005) and lower survival in the larvae (Barker, 1973).

However, density is a relative term, so the implications of a given density depend on the environment and available resources (Bodenheimer, 1938). Likewise, conflicts between male and female fitness optima can be exacerbated by resource availability (Arnqvist & Rowe, 2005; Rowe & Houle, 1996). Males and females often differ in their resource needs, and different nutritional optima or preferences between the sexes are not uncommon (Maklakov et al., 2008). This is especially true during reproduction, when female fecundity generally depends more on protein to maximize offspring production (Jang & Lee, 2025) than carbohydrates, an energy source more important for males than females during reproduction (De Lisle, 2023).

Density dependence can be describes as the number of individuals that can be maintained by a population restricted by resources (Kokko & Brooks, 2003; Kokko & Rankin, 2006). Sex-specific density dependence relates to how population dynamics are effected differently by male vs. female density, so for example if density of one sex has a greater impact on recruitment than another (De Lisle et al., 2022; Miller & Inouye, 2011). How density of each sex influences birth rates in a two-sex population has been modelled as so-called ‘birth functions’ (Miller & Inouye, 2011). As mentioned above, the simplest and most convenient function is female demographic dominance (*N*_*t*+1_ ∝ *F*_*t*_) were only female density matters for offspring recruitment. Other birth functions that incorporate potential contribution of males, have been proposed, namely the harmonic mean 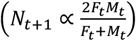 and the geometric mean 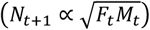 (Caswell & Weeks, 1986). Both harmonic and geometric means uses male and female density, but the harmonic mean is more sensitive to low values in the limiting sex than the proportionate geometric mean is. However, these functions have rarely been empirically evaluated outside of a single study in beetles by Miller & Inouye (2011).

Here, we take an experimental approach with *Drosophila melanogaster* to test the demographic effects of density, sex ratio and adult nutritional environment (high yeast or sugar) on the number of offspring recruited (both total recruitment and per capita recruitment). *Drosophila melanogaster* females and males have different nutritional optima, females require protein for egg production while males need more carbohydrate (Jang & Lee, 2025). Thus, we expect that an adult diet rich in yeast will increase offspring recruitment in comparison with a high sugar diet. While we expect female density to be positively associated with offspring recruitment (see Barker, 1973; Pearl & Parker, 1922; Wallace, 1974), the effects of sex ratio across a density gradient are unclear (Kokko & Rankin, 2006), and provide the opportunity to directly assess candidate models of two-sex population demography. Leveraging experimental population demography in a model system for the study of sexual conflict allows a direct test of how male and female density interact to contribute to population growth in a discrete time system.

## Method

### Data collection

To test the demographic effects males may have on offspring production, we used virgin LHm flies and manipulated sex ratio, density and nutritional environment. The LHm lineage of *D. melanogaster* has been maintained at large population size under a density-controlled, discrete-time life cycle at consistent environmental conditions since collection in California in the 1990s (Rice et al., 2005). This experiment was carried out in a blocked generation design between November 2023 and February 2024 and the experimental flies were reared at 25°C, 60% humidity and a 12:12 (L:D) light regime with normal cornmeal/yeast/molasses-based food and controlled egg density after 18 hours of egg laying. The experimental assay was performed at room temperature under natural light, and the flies were lightly anesthetized with humidified CO_2_ during collection and set up.

Virgins were collected over 1-3 days, always within 3 h of eclosion, until enough flies were obtained to set up one block of the experiment. During collection the virgins were grouped in single-sex vials of 20 individuals and to ensure virginity all female vials that had fertilized eggs before setup were eliminated. To prevent food intake while avoiding desiccation we held the virgins in vials filled with agar (0.017 g/l) until the experiment started.

During the experimental setup each vial (25x95 mm, VWR, Drosophila vials, Narrow style) received one 5µl microcapillary tube (Hirshmann, minicaps, ISO 7550, R≤0.5%, CV≤1.0%) containing either a deionized water solution of 8:1g sugar:yeast (yeast=0.05mg/5µl) or 1:8g sugar:yeast (yeast=0.4mg/5µl) (Table sugar and Yeast extract, bacteriological 1 kg, VWR Life science) per dl (0.09 g/ml) in addition to normal larval food medium. These diet levels were chosen because they reflect high sugar/yeast diets that correspond to increased male/female fitness (De Lisle, 2024). Each vial also received a density of either 2, 4, 10 or 30 flies and a sex ratio between 10 and 90% of each sex (Fig. 1). The flies were then allowed to mate and interact freely for 4 days, because females are most fecund around day 4 (Ashburner et al., 2005) and it’s also appropriate based the LHm culture cycle. A limitation of our diet manipulation is that adults had access to an unlimited amount of larval food media (in addition to the liquid food manipulation) during the experiment; this limitation likely affected our power to detect diet effects, elaborated below.

**Figure 1.**
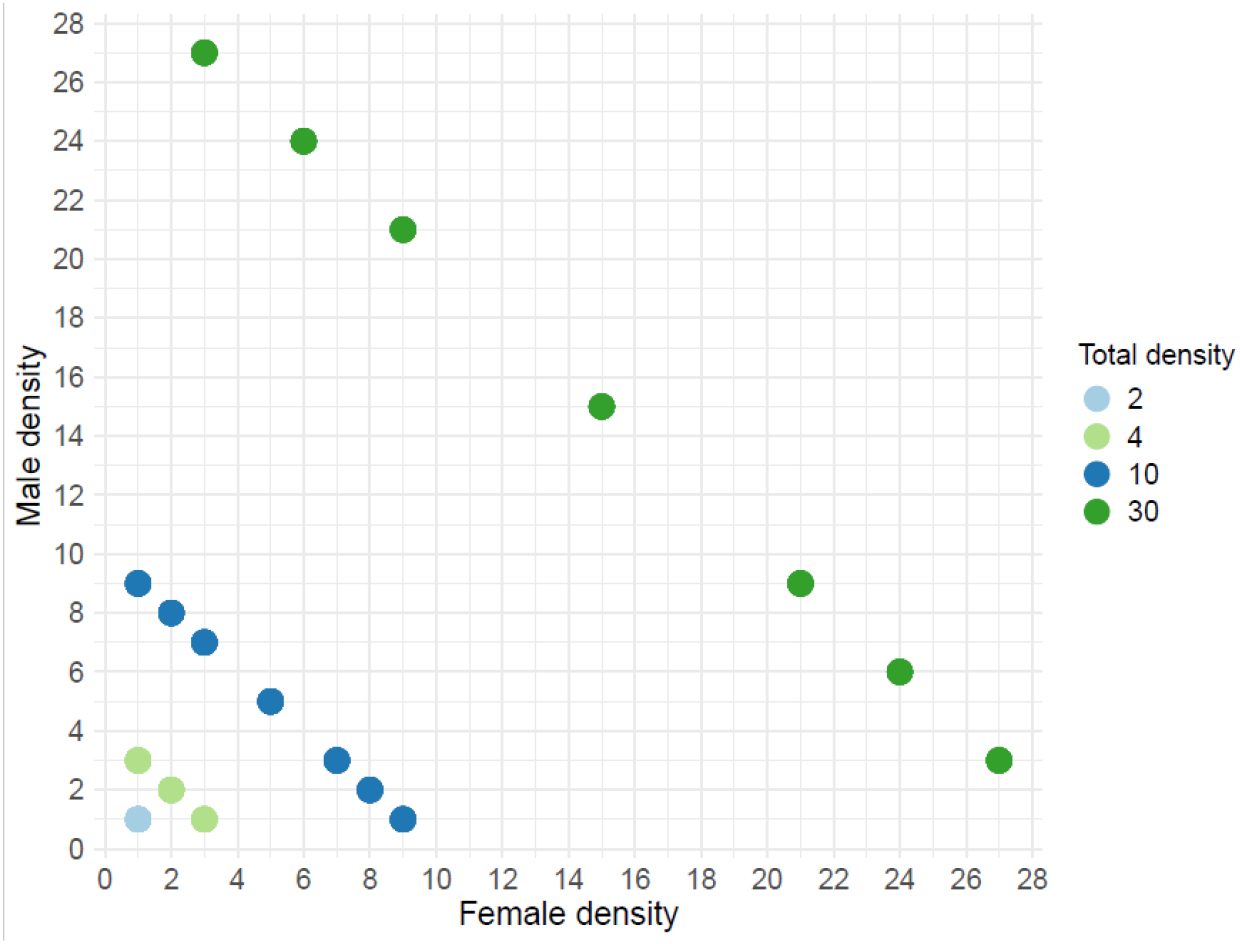
Experiment set up with female and male density. We did the experiment in 5 blocks, 18x2 vials, because of the number of virgins (588) needed for each replicate. All blocks had both food treatments (high sugar vs high yeast) resulting in a total of 2 600 parental flies used including the extra 1:1.

The eclosed offspring were frozen in agar vials 13 days later to be sexed and counted at a later date when we also noted the mean number of offspring per female (hereafter referred to as per capita recruitment). We repeated the experiment across 5 blocks and added some extra replicates of the 1M:1F level in each block to account for low absolute number of adults. One block (B) was missing the highest density (30) due to a shortage of virgins. One vial was eliminated from the analysis due to incorrect data entering.

### Data analysis

Because total offspring recruitment was not normally distributed, we chose a negative binominal generalized mixed model for the analysis of raw total recruitment. For analysis of per capita recruitment, a calculated mean based on female density in the vial, we used a linear mixed effects model. We found no significant effects of our diet treatment on either response variable, and so for simplicity, statistical power, and ease of interpretation, we proceeded by fitting and interpreting models without a diet treatment term, for transparency we report the results below. For each response variable, we fit two different models differing in fixed effect structure. First, we fit a model with sex ratio and total adult density as fixed effects, and a second model with (centred) female and male density as fixed effects. Block was included as a random effect in all analyses. We chose to fit models with sex ratio + density and male + female density because, although they both describe the same information about the treatment levels, they differ in their interpretability. For example, the model with male and female density as fixed effects and per capita recruitment as the response allows an assessment of male contribution to density dependence while the sex ratio and density model shows the bigger picture (sex ratio matters for recruitment).

We also fitted a series of candidate two-sex demographic models to our data using nonlinear least squares. Our candidate models were selected to reflect birth functions hypothesised to characterise demographics of sexual populations. For each birth function (harmonic mean, geometric mean, minimum sex, and female dominance) we fitted four alternative forms of density dependence: density independence, density dependence based on total N (M + F), based on a single sex (F), or two-sex (separate terms for males and females). We proceeded with a Beverton-Holt form of density dependence, which generally fitted the data better than the Ricker model (average delta AIC = 25) (Miller & Inouye, 2011).

The analyses were performed in R (v4.4.2; R Core Team 2024), with the packages “plyr” (Wickham, 2011), “lme4” (Bates et al., 2015), “nlme” (v3.1-166; R Core Team 2024), and the diagrams using “ggplot2” (Wickhem, 2016) and “ggbeeswarm” (v0.7.2; Clarke, Sherrill-Mix and Dawson 2023).

## Results

A total of 20 200 offspring from 191 vials were recorded over the five blocks. No vial was empty but the variation in offspring recruitment was substantial, ranging from 2 to 250 per vial (mean±sd; 105.76±55.48) in total recruitment and between 1.79 and 112 (30.85±23.13) in per capita recruitment. The offspring sex ratio also varied from 0-100% females (52±1) but the actual recruitment was consistent between the sexes with 1-128 sons (51.43±28.93) and 0-131 daughters (54.32±28.22).

### Food treatment

We did not find an effect of the diet treatment on total offspring recruitment (estimate = 3.06, se=7.73, t_185_=0.396, p=0.69, Fig. 2A), per capita recruitment (estimate = 2.37, se=3.19, t_185_= 0.74, p=0.46, Fig. 2B) nor offspring sex ratio (estimate =0.003, se=0.014, t_185_=0.18, p=0.86, Fig. 2C) but the estimates implies that yeast increase recruitment. Neither did a Levene’s test show a difference between the variation within diets (total recruitment, F=2.15, p=0.14; capita recruitment, F=0.315, p=0.575; offspring sex ratio, F=1, p=0.319).

**Figure 2.**
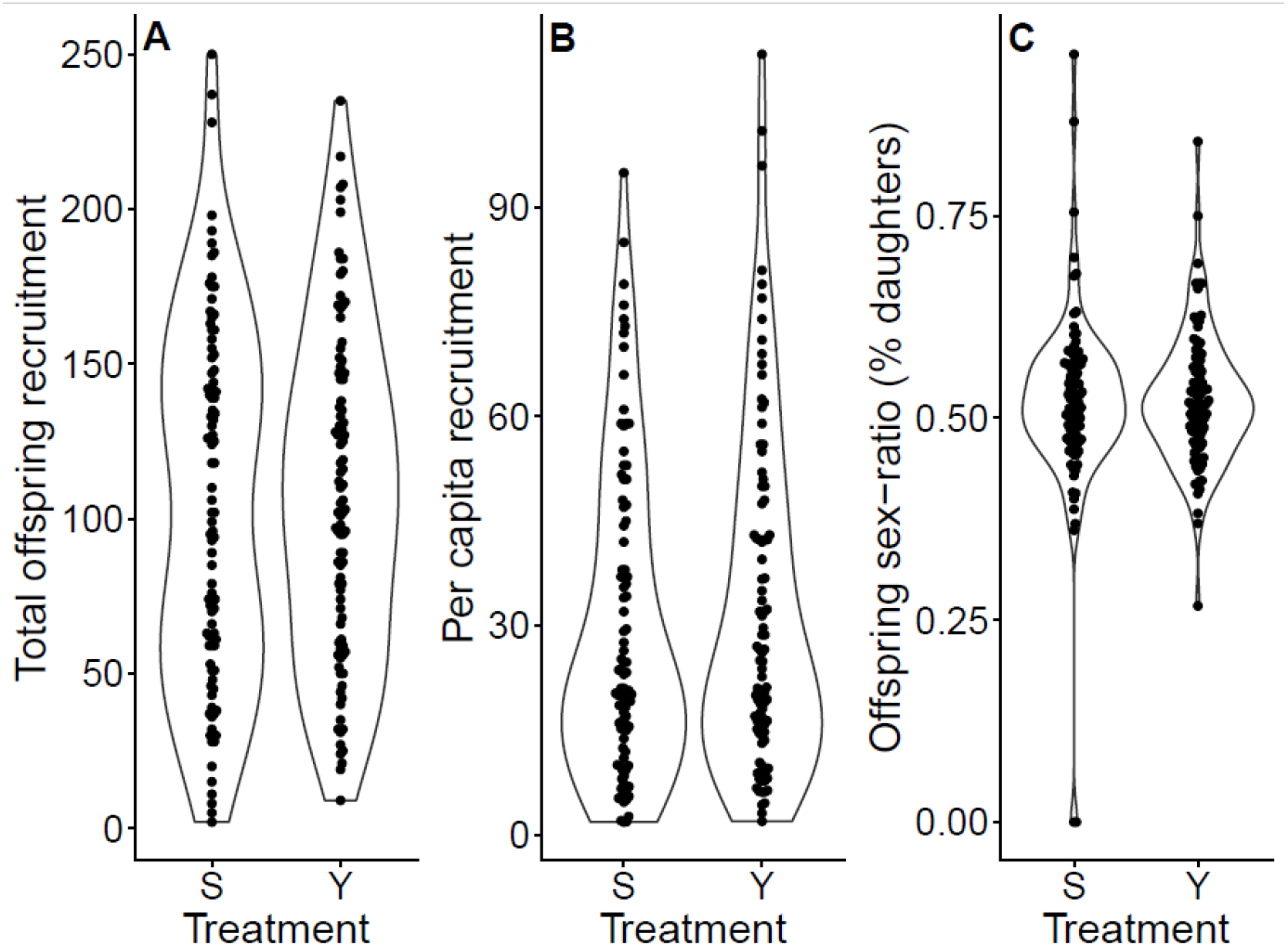
Non-significant effects of the high sugar (8:1 g/dl) (n=95) and yeast (1:8 g/dl) (n=96) treatment on A) total offspring recruitment, B) per capita recruitment and C) offspring sex ratio.

### Total recruitment

When analysing density and sex ratio as predictors, we found that density (estimate =0.03, se=0.008, z_185_=4.5, p=5.54^e-06^), sex ratio (estimate =1.87, se=0.27, z_185_=6.77, p=1.23^e-11^) and the interaction between them (estimate =-0.04, se=0.01, z_185_=-2.62, p=0.008) had an effect on total offspring recruitment (Fig. 3A). A similar pattern is shown when analysing the male and female density where we saw that only females had an effect (estimate = 0.04, se= 0.006, z_185_=7.44, p=1^e-13^) on total offspring recruitment, the male density (estimate =-0.005, se=0.005, z_185_=-0.91, p=0.361) and the interaction between female and male density (estimate = 0.002, se=0.001, z_185_=1.34, p=0.18) were not statistically significant predictors of total offspring recruitment. These results show that more females, both in percent and actual numbers, produce more offspring regardless of the male density (Fig. 3A). We also see that when the sex ratio reaches 50% females and above, there is no difference between density 10 and 30 flies per vial in terms of total offspring produced, while for male-biased sex ratios at a density of 30 adults produces more offspring than at a density of 10.

**Figure 3.**
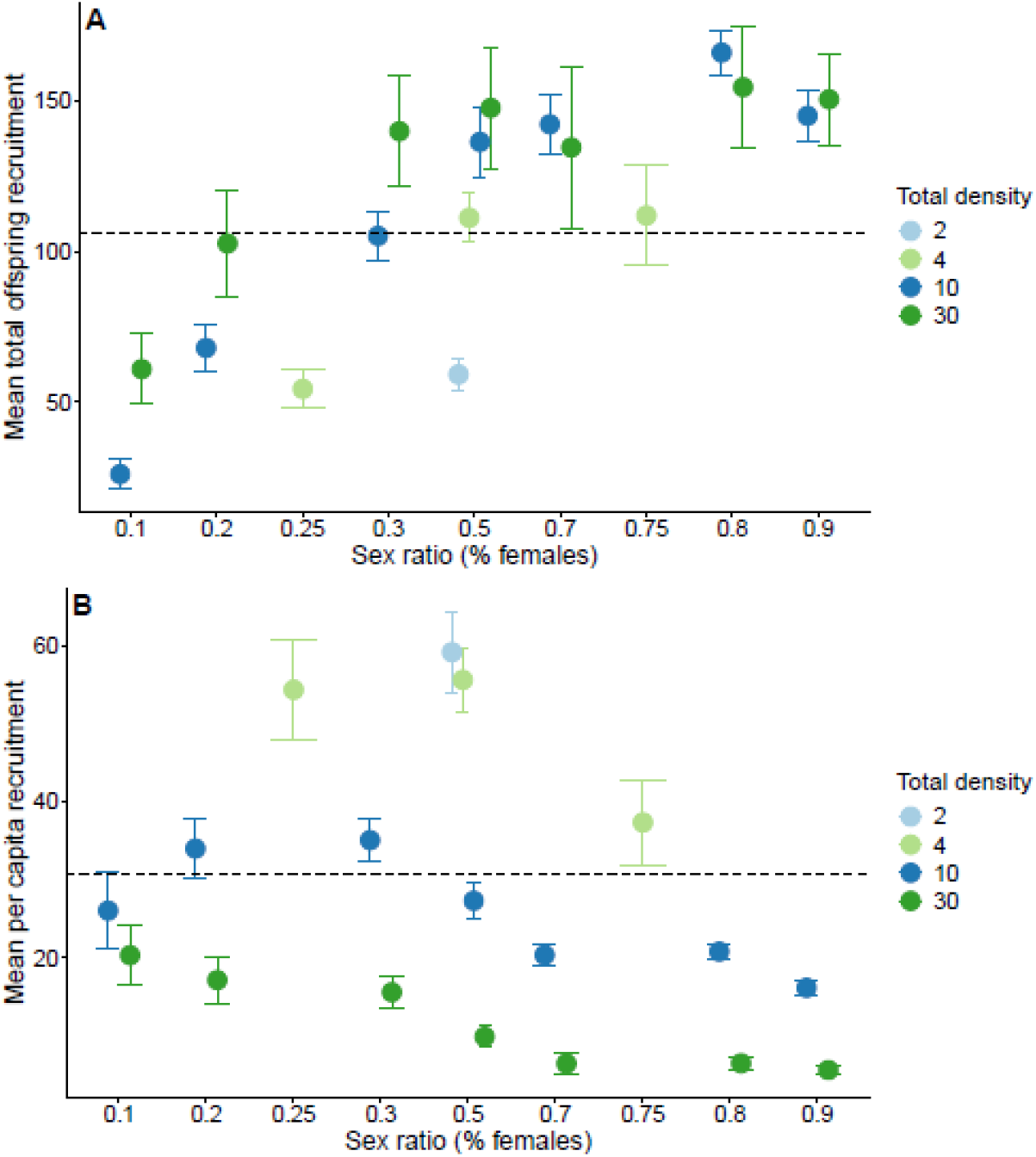
Mean values from the five blocks stating the relationship between sex ratio, total density and female density in the parental generation and A) total offspring recruitment, B) per capita recruitment. 2 600 parental flies resulted in 20 200 offspring which gives a mean recruitment of 105.76 (dashed line A) and mean per capita recruitment of 30.8 (dashed line B) per female for the total experiment. Error bars=±1 SE

### Per capita recruitment

The analysis of density and sex ratio reveal both density (estimate =-1.41, se=0.25, t_183_=-5.67, p=0.000) and sex ratio (estimate =-24.12, se=9.05, t_183_=-2.67, p=0.008) as significant predictors of per capita recruitment, but the interaction (estimate =0.21, se=0.45, t_183_=0.46, p=0.64) was not significant (Fig. 3B). In a model with male and female density as predictors, both male density (estimate=-0.81, se=0.17, t_183_=-4.85, p=0.00) and female density (estimate=-1.63, se=0.17, t_183_=-9.69, p=0.00) were significant, with females having nearly twice as large effect size as males (-1.63 vs -0.81). This model also revealed a significant interaction between male and female density (estimate = 0.09, se=0.04, t_183_=2.37, p=0.019) on per capita recruitment. This means that each female has higher fitness when the competition from both females and males is lower (Fig. 3B), and shows that low densities with fewer females than males result in higher per capita recruitment. However, we found that when female density is low (1-3) females at equal sex ratios, regardless of density, have a higher per capita recruitment, a pattern lost at higher female densities ≥9. Likewise, our results show that density has a more pronounced effect when sex ratios are 20% or higher, at 10% (a strongly male-biased sex ratio) a smaller difference between density levels can be seen (Fig. 3B). These results indicate that male and female density both can affect per capita recruitment, but density dependence is far stronger when acting through females than when acting though males.

### Birth functions

We fitted candidate demographic models to our data and found that the two-sex density dependence (where both male and female density contribute to density dependence in the form of 1/(b_M_*M + b_F_*F)) always fits better (based on AIC) than one-sex (where only females contribute to density dependence, 1/ b_F_*F), total density (1/(b*(M+F)) and density independence (Table 1). The best fitting model, by 16 AIC units, was a female dominance birth function with sex-specific density dependence (Table 1). Together, our results indicate that females matter most for offspring production, but males have a role in contributing to density dependence (Table 2, Fig. 4), although parameter estimates indicate that male contribution to density dependence was substantially weaker than that of females (Table 2, Fig. 4), consistent with the results from the experiments on offspring recruitment.

**Table 1.** Comparison of the fitted Beverton-Holt birth functions.

| Birth Function | Density Dependence | No. Parameters | $\Delta AIC$ |
| --- | --- | --- | --- |
| Female Dominance | Two-Sex Beverton-Holt | 3 | 0.00 |
|  | One-Sex Beverton-Holt | 2 | 16.70 |
|  | Total Beverton-Holt | 2 | 83.57 |
|  | None | 1 | 253.21 |
| Harmonic Mean | Two-Sex Beverton-Holt | 3 | 38.31 |
|  | One-Sex Beverton-Holt | 2 | 231.40 |
|  | Total Beverton-Holt | 2 | 112.48 |
|  | None | 1 | 243.56 |
| Geometric Mean | Two-Sex Beverton-Holt | 3 | 16.58 |
|  | One-Sex Beverton-Holt | 2 | 233.00 |
|  | Total Beverton-Holt | 2 | 86.89 |
|  | None | 1 | 235.62 |
| Minimum | Two-Sex Beverton-Holt | 3 | 82.86 |
|  | One-Sex Beverton-Holt | 2 | 234.38 |
|  | Total Beverton-Holt | 2 | 171.84 |
|  | None | 1 | 265.68 |

**Table 2.** Beverton-Holt best fitting birth function, Two-sex female dominance.

Parameter estimates and standard errors for best-fit demographic model
| Parameter | Estimate | Std. Error | t valuep value |
| --- | --- | --- | --- |
| $\lambda$ | 115.9457314 | 17.7645914 | 6.526789<0.001 |
| bF | 0.6547280 | 0.1175990 | 5.567463<0.001 |
| bM | 0.0828036 | 0.0311007 | 2.6624330.008 |

**Figure 4.**
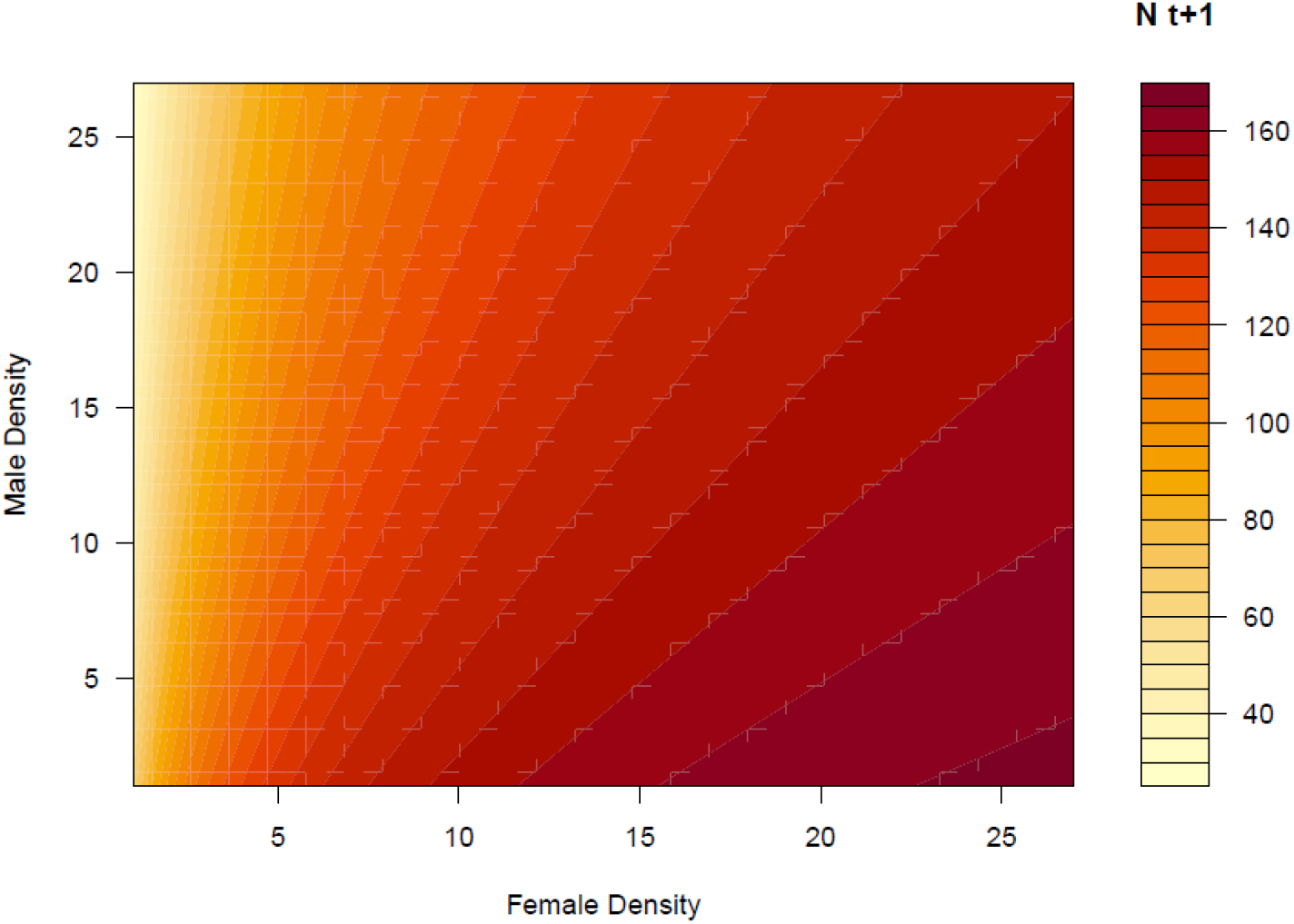
Figure shows predicted values of population size at the next generation as a function of male and female density, from best-fitting demographic model (see Table 1, 2).

## Discussion

We performed an experimental manipulation of sex ratio and density across two food treatments to assess the demographic effects male and female density had on offspring production in *D. melanogaster*. This experiment allowed us to understand how a common assumption of population demographic models may be violated in a system where A) the population is adapted to discrete time generations and B) sexual conflict is pervasive. We measured total offspring recruitment, including sex ratio in the offspring, and per capita recruitment. We found that total density, sex ratio and female density matter significantly for both per capita recruitment and total offspring recruitment, but in opposite directions, as expected in the presence of density dependence under reasonable density ranges. Female-biased sex ratios resulted in more offspring than male-biased sex ratios, but each female produced fewer offspring. Our food treatment did not affect the offspring recruitment, neither total recruitment nor per capita recruitment or the offspring sex ratio. The most important contribution of our work is demonstrating a significant role for males in contributing to negative density dependence in a system otherwise characterized by female demographic dominance.

We found a positive effect of sex ratio (increased male-biased sex ratio) on per capita recruitment. This can be explained by decreased larval competition (Barker, 1973) and weaker density dependence via adult males versus females. That is, females are better off (in terms of producing offspring successfully recruited to the next generation) at higher male instead of female densities but would be even better off without the surplus males. However, at the three most male-biased sex ratios at the three lowest densities each female performs better when the sex ratio is approaching 50%, this is probably due to sexual conflict because when the female density is 9 or higher, we no longer see this effect. We see the same pattern with total offspring recruitment, more equal sex ratio at lower densities results in more offspring. Higher female sex ratios may result in more offspring recruited in total for the obvious reason that there are more females to lay eggs, but likely also because mating is known to harm female *Drosophila* (Lew et al., 2006; Pitnick & García–González, 2002) and at higher female densities each female may mate fewer times than at male-biased sex ratios (Rankin et al., 2011). Even so the most male-biased sex ratios may have a reduced mating probability due to high male-male competition and interference (Gomez-Llano et al., 2018).

Unsurprisingly, we observed opposite results when analysing per capita recruitment and total offspring recruitment, which is consistent with those of Wallace (1974) and Pearl & Parker (1922), namely that higher female density results in more offspring but lower per capita recruitment. This pattern could emerge because at low density, larvae competition decreases, and fecundity is limited by sexual conflict (Arnqvist & Rowe, 2005; Barker, 1973) rather than density dependence due to larval competition. However, when density increases, stronger larval competition decreases female per capita recruitment (Barker, 1973). Larval mortality is important and plays a crucial role in population dynamics (Bodenheimer, 1938; Chiang & Hodson, 1950; Sang, 1949). High larvae density can affect survival, but if the larvae die early the competition among surviving larvae will not increase (Miller, 1964). Likewise, more eggs laid also contribute to higher larvae density, which could be counterproductive if higher densities lead to higher larvae mortality. Also, Pearl (1932) tested if higher densities resulted in a lower egg production for each female and concluded that was the case. We did not count the eggs, but it is reasonable to think that our flies also had a lower fecundity at higher densities. There is a point when adding more females will decline offspring recruitment (Barker, 1973), but we did not reach that point in this experiment.

No effect of nutritional environment was found, which was surprising given that yeast intake is well known to positively influence female fecundity in flies (Ashburner et al., 2005). The lack of food effect could be due to too little added liquid food media (5µl per vial regardless of adult density), contributing to increased competition instead of being a resource obtainable by most individuals. Also, because standard fly food was available in each vial in addition to the liquid food media during the entire 4 days of reproduction, it is likely this shared nutritional resource contributed to weaking any effect on our food manipulation. Even though the protein in standard fly food is likely to be relatively inaccessible to adult flies, the accessible sugar and the nonlimited access to food overall is likely to be a confounding factor due to making yeast-abundant levels virtually non-existing. Thus, it is likely that a combination of high larvae density and adult access to larval food limited our ability to assess potential nutritional effects on demographic parameters. Future experiments that manipulate adult resources independently of larval food availability may be informative.

We also fitted a set of candidate birth functions to our data, linking our experiment to theoretical models of two-sex population demography (Miller & Inouye, 2011). Our finding of strong support for female demographic dominance with sex-specific negative density dependence is consistent with the results of our linear models regarding total offspring recruitment, namely that females play the greatest role in both “births” but also in contribution to negative density dependence. In this case males matter primarily through their effects on density dependence, and maybe this is generally true in systems like flies where there is widespread sexual conflict, including harassment, and every female can produce large clutches (Chapman et al., 2003). A noteworthy conclusion of our analysis is that, while increasing male density has negative effects on female per capita recruitment, consistent with a substantial body of work indicating costs of male mating behaviour for female fitness (Arnqvist & Rowe, 2005), the effects of increasing male density are substantially weaker than the effect of increasing female density.

In population demography males can matter in different ways depending on their role in offspring production (Arnqvist & Rowe, 2005; Rowe & Rundle, 2021). Most demographic models see males as only sperm donor, but males may often influence population dynamics in more direct ways (Caswell & Weeks, 1986). Our results indicate that male harassment can lead to males contributing directly to density dependence. Yet, male density dependence is about half the effect size compared to females; female density still has a far stronger negative effect on per capita recruitment than males. This puts the concept of sexual conflict in a demographic framing and points to males mattering for offspring recruitment and thus population dynamics, at least to some extent. Our results indicate that males matter less when the total density is very low, or when sex ratio is strongly female-biased. Thus, males “matter” here not because of allee effects when rare (as is sometimes proposed as a justification for two-sex demographic models (De Lisle et al., 2022)), but rather because of sexual conflict when densities are high. From these results we can draw the conclusion that even if a population can be characterized as female demographic dominant, it doesn’t mean that males are unimportant. Especially for conservation and wildlife management it would be a good idea to investigate how or if both males and females contribute to population dynamics before focusing the effort to only on one sex.

Remarkably few studies have taken an experimental approach to evaluate candidate models of two-sex population demography (but see Miller & Inouye, 2011). Hence the importance of our study which combines several of the natural occurring variables in a population, not only in *Drosophila* but in many species. In order to better understand how population demography affect offspring recruitment and population dynamics, future experiments should use larger populations with more extreme sex ratios to reach the point where higher female densities lower the total amount of offspring recruited.

## Acknowledgment

We thank Folmer Bokma and Ted Morrow for comments on earlier drafts of the manuscript. Founding was provided by grants from the Swedish Research Council (Vetenskapsrådet: grants no. 2019-03706 and 2024-04599), and KK-stiftelsen (grant no. 2024STG).

## Authors’ contributions

All authors contributed to designing and performing the experiment, A.B collected data, performed the analysis, and drafted the manuscript, all authors contributed to revising the manuscript.

